# Evolution towards increasing complexity through functional diversification in a protocell model of the RNA world

**DOI:** 10.1101/2021.05.10.443418

**Authors:** Suvam Roy, Supratim Sengupta

## Abstract

The encapsulation of genetic material inside compartments together with the creation and sustenance of functionally diverse internal components are likely to have been key steps in the formation of ‘live’, replicating protocells in an RNA world. Several experiments have shown that RNA encapsulated inside lipid vesicles can lead to vesicular growth and division through physical processes alone. Replication of RNA inside such vesicles can produce a large number of RNA strands. Yet, the impact of such replication processes on the emergence of the first ribozymes inside such protocells and on the subsequent evolution of the protocell population remains an open question. In this paper, we present a model for the evolution of protocells with functionally diverse ribozymes. Distinct ribozymes can be created with small probabilities during the error-prone RNA replication process via the rolling circle mechanism. We identify the conditions that can synergistically enhance the number of different ribozymes inside a protocell and allow functionally diverse protocells containing multiple ribozymes to dominate the population. Our work demonstrates the existence of an effective pathway towards increasing complexity of protocells that might have eventually led to the origin of life in an RNA world.

## 1. Introduction

The RNA world hypothesis posits a central role for RNA in information storage, catalysis and regulation and argues that life based on RNA only existed prior to a DNA-protein world. The discovery of ribozymes [1–3], their synthesis in the laboratory using *in vitro* evolution [4–6] and abiotic synthesis of ribonucleotides [7–11] have provided indirect evidence for the plausibility of an RNA world. Several theoretical models have complemented the exploration of the pure RNA-world scenario in the laboratory. They provide further proof of the plausibility of life based on RNA sequences and identify the conditions under which such functional RNA sequences can emerge via non-enzymatic processes. The thermal gradients produced near hydro-thermal vents have been shown to be useful in monomer accumulation and ligation, therefore paving the way for emergence of complex RNA replicators [12, 13]. Wet-dry cycles have also been shown to be effective for formation of long RNA polymers [14, 15]. Template-directed ligation with a limited pool of monomers were shown to lead to the formation of RNA strands with high compositional diversity [16]. In a previous work we have studied the earliest epochs of the RNA world that eventually lead to the emergence of functional RNA sequences like ribozymes and tRNA-like structures from basic monomer building blocks through non-enzymatic, template-directed primer extension and concatenation reactions.

Recent research suggests that it is likely that RNA instead of being exclusive players in a primordial world, coexisted with small peptides [17, 18] and lipids [19, 20]. Lipid vesicles, that may have formed through self-aggregation of hydrophobic lipid molecules to form closed compartments [21, 22], can provide several advantages [23] in an RNA world. The co-existence of RNA sequences, short peptides and lipids opens up the possibility of compartmentalization of RNA sequences in lipid vesicles which could potentially grow [24] due to transfer of material from other empty oleate vesicles [25, 26]. The type of vesicular membrane that facilitate non-enzymatic replication of RNA strands [27] and catalytic action of encapsulated Hammerhead ribozymes [28] have also been identified. Such experiments are gradually revealing the conditions under which vesicles can start acting as chemical factories that eventually triggered the transition from chemistry to biology and lead to the emergence of the the earliest protocellular life-forms. Computer simulations have further extended the insights obtained from chemical experiments. They have revealed that encapsulation inside protocells leads to selection of replicases [29]. RNA replicators have been shown to have enhanced likelihood of survival inside lipid vesicles when compared to surface-based, spatially open systems [30, 31] because compartments provide more effective protection against hydrolysis and take-over by parasitic counterparts. Protocells also have the advantage of enhancing the maximum error-threshold [31] that can be tolerated during replication of ribozymes, thereby making such functional sequences more robust against mutational degradation. Allowing horizontal transfer of molecules between protocells can further stabilize the replication of fragmented ribozymes inside them [32].

Despite these manifold advantages of protocells, there are several questions that need to be answered before we can fully understand how RNA-based life encapsulated within protocells emerged. The emergence of the first ribozyme inside a protocell must have been a chance event. Even if the template-directed replication process produces a large number of replicates, how likely is it that one of them become a functional molecule like a ribozyme? Although a precise answer to this question remains elusive, recent work [33–35] suggest that this number may not be too low. Random sampling of sequences reveal that only a small set of secondary structures are obtained upon folding the sampled sequences with many sequences yielding the same secondary structure. Certain secondary structures appear much more frequently and such strong phenotype bias is evident from the large (several orders of magnitude) differences in the frequencies of the most and least common secondary structures. Remarkably, the secondary structures found in nature happen to be the ones that appear with the highest frequency [35]. Moreover, the number of random sequences that need to be sampled to generate these secondary structures is quite low (*≈*10^5^ for L=126) compared to the total number of possible sequences (4^*L*^) which can be astronomically large for large L [35]. These results suggest that functional structures may not be very difficult to produce and can be generated by sampling a relatively small number of RNA sequences. Nevertheless, we need a better understanding of the processes of RNA replication that favoured the appearance and subsequent proliferation of the first ribozyme. In other words, what were the processes by which the first ribozyme came into existence inside the protocells? What was its functional nature and what impact did it have on protocellular evolution ? If subsequently, other ribozymes appeared during the error-prone replication process, were protocells with multiple ribozymes having diverse functionality able to proliferate at the expense of those with fewer ones?

In this work, we use the rolling-circle mechanism of RNA sequence replication within protocells to address these questions. We show that this mechanism, first observed in viruses [36– 38], is more effective than the non-enzymatic template-directed primer extension process in creating a large number of replicated RNA sequences. The chance creation of a replicase ribozyme with a small probability during the error-prone RNA sequence replication process inside protocells can further enhance the RNA replication rate. If other functionally distinct ribozymes also appear subsequently, under certain conditions, they can act synergistically to enhance each other’s production. We speculate on the nature and hierarchy of the emergent ribozymes and show that they can not only synergistically aid each other’s formation but also favour evolution of the protocell population towards increasing complexity through preferential selection of protocells with larger number of functionally diverse ribozymes. Our work shows how the rolling circle mechanism of replication together with the chance creation of a few ribozymes can work in conjunction and point to a plausible pathway for the emergence of the earliest protocellular life-form.

## 2. Material and methods

A replication mechanism that can efficiently create a large number of RNA sequences is more likely to yield a ribozyme as the outcome of a chance replication event. We compared RNA replication by the non-enzymatic, template-directed primer extension mechanism with the rolling circle mechanism to determine their relative effectiveness. These comparison simulations (see *Supplementary material* for details) reveal that exponential growth of the number of strands necessary to sustain the replication process against degradation in case of template-directed primer extension, is constrained by the requirement of very high dry-phase temperatures *≥* 60° C [Fig-S1]. It therefore seems likely that for more realistic dry phase temperatures that were prevalent in primordial earth, the template-directed primer extension process may not have been effective in sustaining high growth rate of RNA strands. In contrast, the rolling circle mechanism can lead to exponential growth of the number of both open-ended single-stranded RNA (ssRNA) and circular double-stranded RNA (dsRNA) molecules [Fig-S2] and is not constrained by the requirement of a dry phase (see *Supplementary material* for details). We therefore consider the creation of new RNA sequences inside protocells to be driven by the rolling circle mechanism.

### (a) Rolling circle mechanism of RNA replication

We briefly describe the rolling circle replication mechanism. In this process (see Fig-1) a small complementary primer (*∼* 8 nt) first attaches to a circular ssRNA. The primer then extends by template-directed primer extension. Upon becoming full length it can extend further by gradually displacing its other end from the initial point. This creates a hanging tail which keeps growing in length with further primer extension. However when the hanging tail becomes too long, it breaks off and becomes an open-ended ssRNA.

**Fig. 1.**
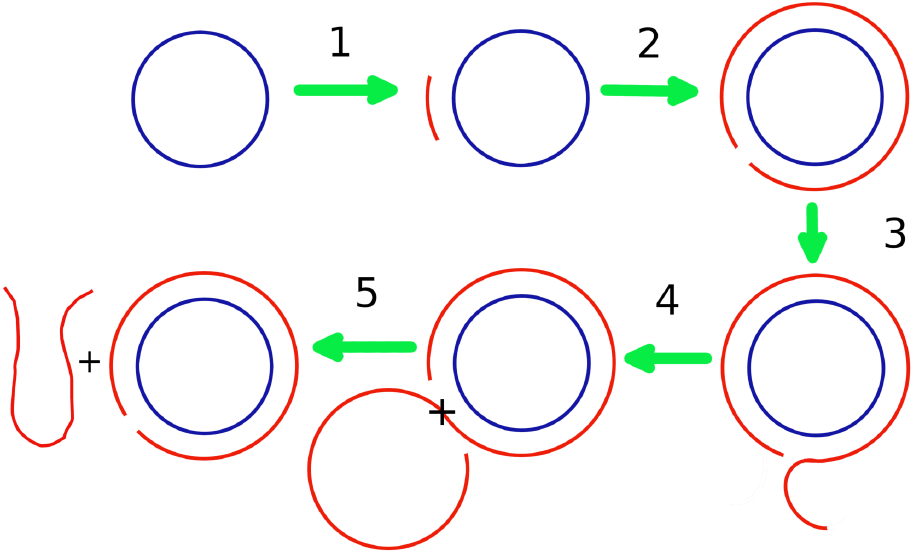
Schematic diagram of the rolling circle replication process. **1:** A short complementary primer attaches to a circular ssRNA template. **2:** It is extended by the template-directed primer extension process towards full length. **3:** Upon attaining full length, the primer extends further by displacing the other end of itself from the initial point, resulting in an overhanging portion. **4:** when the overhanging tail attains a length equal to that of the circular template, it breaks apart. **5:** The separated tail becomes an open-ended ssRNA.

### (b) Protocell model

We start with a model consisting of N lipid vesicles (protocells) each of which encapsulate one circular ssRNA molecule of length 200 nucleotides [39]. Reactions occurring inside each protocell (described below) with specified rates leads to formation of different species and quantity of RNA molecules. The evolution of the protocell population occurs following a birth-death process, when a protocell containing a number of RNA species (*V*_*i*_) that is greater than a specified threshold (*V*_*T*_) splits into two with its contents divided at random between the daughter cells. To keep the population size fixed, a protocell (j) is eliminated with a probability that is proportional to the size difference (*V*_*i*_ *−V*_*j*_). This ensures that protocells with larger number of RNA strands are more likely to survive across generations. However, our results do not change significantly even if the eliminated protocell is selected at random (see Fig-S3(A)).

Initially when the protocells contain only circular RNA, three types of reactions are possible: conversion of circular ssRNA (s) to circular dsRNA (d) via template-directed primer extension process, production of new open-ended ssRNA (l) from circular dsRNA (d) through the rolling circle replication mechanism and degradation of all types of RNA strands. The ssRNA molecules produced through rolling circle replication will mostly fold into complex structures because of their long lengths. Due to mismatches during replication, the replicated strands will not be exact compliments of the template. We assume that a small fraction of those folded ssRNA molecules with complex structures attains enzymatic capabilities [15] like that of a replicase (r), cyclase (c), nucleotide synthase (n) and peptidyl transferase (p). This expectation is based on the fact that error-prone replication of the same template at different times can produce sequences with distinct, complex secondary structures. Fig-S4 in the Supplementary material shows a subset of distinct secondary structures of sequences obtained by non-enzymatic replication of the same 200-mer template sequence multiple times. Such structural diversity at 200 nt length is not surprising as Table-S1 shows the increase in structural diversity with increasing template lengths. Given their complex secondary structures, as well as for reasons given in the previous section, it may not be unreasonable [33– 35] to expect that some of these replicates can exhibit distinct catalytic abilities.

A replicase can catalyze the process of circular ssRNA to circular dsRNA and circular dsRNA to open-ended ssRNA production by increasing the replication rate to *K*_*fast*_ *>> K*_*rep*_ which is the non-enzymatic replication rate. We use *K*_*fast*_ *∼*0.362 *h*^*−*1^, which is the rate at which the fastest replicase found [6] can replicate a 200-mer. The process of replicase catalyzed replication though significantly fast, is still prone to mismatches which increase sequence diversity in the protocell. Hence, we assume that new ribozymes can be created with same probabilities even in such cases. A cyclase can circularize an open-ended ssRNA molecule to form a circular ssRNA by joining its open ends. As there are no experimental rates available for the circularization reaction, we assume this also occurs at the rate *K*_*cyc*_ = *K*_*fast*_. A nucleotide synthase can create new monomers inside a protocell in situations where the finite monomer pool is inadequate for all the reactions to proceed and is therefore crucial for sustaining new RNA strand formation in monomer-poor environments. Finally a peptidyl transferase (p) can join free amino acid molecules to form small peptide chains [40]. Lipid molecules of the protocell membrane turn lipophilic by forming compounds with small peptide chains [25] and such lipophilic membranes attract lipid molecules from nearby protocells with non-lipophilic membranes and grow at the expense of the latter. Hence, we assume that whenever peptidyl transferase ribozymes appear inside a protocell ‘i’, it will grow in size i.e. its upper threshold volume *V*_*T*_ will increase as 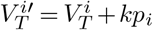 (where k is the amount of volume increase in units of RNA strand numbers brought about by a single peptidyl transferase ribozyme). In the presence of these ribozymes there will be 3 extra reactions namely; replicase catalyzed circular ssRNA to circular dsRNA, circular dsRNA to open-ended ssRNA and cyclase catalyzed, non-catalytic open-ended ssRNA to circular ssRNA creation.

For a finite monomer system we multiply each replication rate with a term 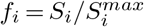 to account for reduction of replication rates in absence of sufficient monomers. Here 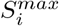 denotes the maximum number of monomers (in units of 200-mers) a cell can hold and *S*_*i*_ is the number of available monomers in a cell at any instant in units of 200-mers. The initial number of free monomers in each protocell was taken to be 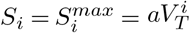, where *a* is a constant factor. In the presence of nucleotide synthase, new monomer creation inside a cell will depend on the number of nucleotide synthase molecules inside that cell. Hence we modify the multiplication factor as 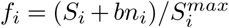. The quantity ‘b’ is a measure of the number of free monomers (in units of 200-mers) a nucleotide synthase can produce. The *S*_*i*_ values after each reaction step are updated to 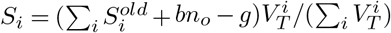, where 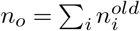 is total number of nucleotide synthase molecules in the entire population before reaction and *g* is the number of new strands created in the entire population after the reaction step.

The reactions described above lead to changes in numbers of different RNA species inside the i’th protocell. The differential equations expressing the variation in the number of each type of RNA species over time are:

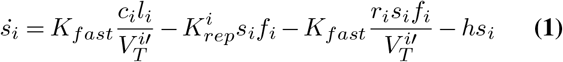

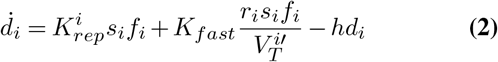

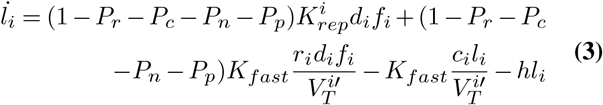

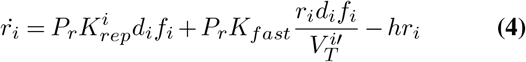

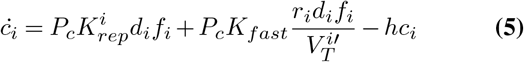

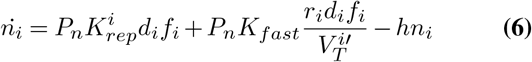

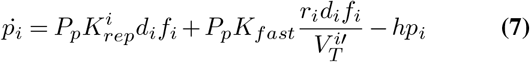

where s, d, l, r, c, n, p denote the number of circular ssRNA & dsRNA, open-ended ssRNA, replicase, cyclase, nucleotide synthase and peptidyl transferase respectively. *P*_*r*_, *P*_*c*_, *P*_*n*_, *P*_*p*_ are the creation probabilities of these 4 types of ribozymes by the rolling circle replication process. The degradation rate is taken to be same (*h* = 0.0008 *h*^*−*1^) for all RNA species. The ribozyme-catalyzed reactions are second order reactions i.e. the formation rate of the corresponding RNA species will be proportional to the number of both ribozymes and substrates. However as the L.H.S. of the rate equations denote the rate of change of the number of different types of RNA species, we divide these second order rates with the maximum protocell volume 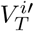 to match the dimensions.

Primers which attach to circular 200 nucleotide long ssRNA templates to start off the process of replication by the rolling circle mechanism have to be at least 8 nt long in order to form a stable pair with the template. Vesicle membranes are known to be permeable to monomers and short oligomers [27, 28]. Here we assume that starting from monomers, oligomers up to 8-mers can penetrate a protocell’s membrane from both direction to ensure the availability of 8-mer primers among all protocells. To account for the difference in sequence of the original circular ssRNA templates in different protocells, the non-enzymatic replication rates of 200-mer circular ssRNA molecules initially present in each protocell were chosen from the distribution 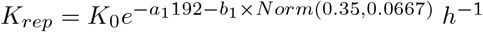 (neglecting 8-mer primer annealing times which are very small compared to replication times). This distribution was obtained from sequence-level simulations that are described in detail in the *Supplementary material*; which discusses how *K*_*rep*_ changes with the relative number of mismatches and strand length (see Fig-S5).

The 6 types of reactions that can occur inside a protocell can be divided into two categories; non-enzymatic: circular ssRNA to circular dsRNA, circular dsRNA to open-ended ssRNA & degradation of all kind of RNA strands and enzymatic: replicase catalyzed circular ssRNA to circular dsRNA, replicase catalyzed circular dsRNA to open-ended ssRNA & cyclase catalyzed openended ssRNA to circular ssRNA. We calculate total reaction rate for each cell and then determine the time step size dt as inverse of the maximum of those rates 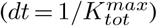. During this time period a cell will undergo any of the 6 types of reactions with probability 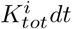. If a reaction occurs, the type of reaction is chosen at random based on the relative reaction propensities of these6 types of reactions, which are 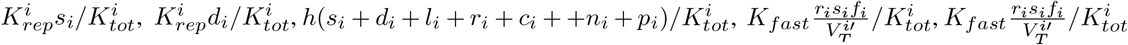 and 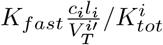 respectively. The cell division and population update steps are also implemented stochastically, as described earlier.

## 3. Results

Even though our ultimate goal is to ascertain the outcome of competition between different protocells in a population that are distinguished by the number and type of their component RNA species, it is instructive to use the above dynamical model to find out the effect of varying probabilities for creation of different types of ribozymes inside a single protocell. We start with a single circular ssRNA molecule inside a protocell with non-enzymatic rate *K*_*rep*_ *∼*0.0082 *h*^*−*1^ and solve the Eqs.1-7 numerically to determine the time evolution of the number of different types of RNA strands inside it. For this numerical model we ignore the process of 8-mer primer attachment to a circular ssRNA. We first consider the case of abundant monomer availability and use *S* = *S*_*max*_ = *aV*_*T*_ = 50*V*_*T*_ and neglected the possible formation of nucleotide synthase, peptidyl transferase (i.e. *P*_*n*_ = *P*_*p*_ = 0). We varied the creation probabilities of replicase (*P*_*r*_) and cyclase (*P*_*c*_) while keeping their sum fixed to unity. Fig-S6(A) shows that a faster rate of growth of RNA strands is achieved when *P*_*c*_ *> P*_*r*_. For a single protocell, the predominance of cyclase over a replicase is more important since the former leads to the creation of many new circular templates which enhances the likelihood for creation of dsRNA and eventually potential ribozymes (both cyclase and replicase). Since the production of ribozymes, including replicases, depend on the availability of circular templates, insufficient number of circular templates on which the replicase can act, creates a bottleneck for production of new ssRNA strands. For a monomer deficient system (*S*_*max*_ = *aV*_*T*_ = 0.8*V*_*T*_), the presence of a nucleotide synthase that catalyzes monomer production can help in sustaining the growth of new RNA strands. Lack of adequate monomers can affect growth of new strands despite the presence of cyclase and replicase and lead to saturation in number of RNA strands. Growth of RNA strands is most favoured when the formation probability of a nucleotide synthase (*P*_*n*_) is similar to that of a cyclase and replicase, *P*_*r*_ = *P*_*c*_ *∼P*_*n*_ [Fig-S6(B)]. And finally we assumed the creation of peptidyl transferase that catalyses formation of short peptides and in the process facilitates protocell growth via membrane transfer. We varied its creation probability (*P*_*p*_) by keeping the sum of all four ribozyme formation probabilities fixed and assuming *P*_*r*_ = *P*_*c*_ = *P*_*n*_. Increasing *P*_*p*_ gradually decreases RNA strand formation rate [Fig-S6(C)] even though it leads to higher threshold volumes. These results suggest a hierarchy for the appearance of ribozymes with the cyclase being the most important in our framework, followed by the replicase. In monomer poor environments, the probability of formation of nucleotide synthase also need to be larger than a threshold (see Fig-S6(B)) to ensure an adequate supply of monomers needed for sustained growth of strands. Even though the peptidyl transferase is helpful in facilitating protocell growth and diversification of its component RNA species, its impact is dependent on the presence of the other enzymes. However, these results which address the evolution of RNA species inside a single protocell, do not reveal whether competition between protocells with different RNA content selectively favour certain types of protocells. We address this issue in the subsequent sub-sections. Intriguingly, we find that the conditions for the proliferation of protocells with increasingly diverse functionality are distinct from the conditions required for the highest rate of growth of RNA strands inside a single protocell.

### (a) Role of replicase and cyclase in protocell evolution

Using our fully stochastic population dynamics model, we examined the effect of replicase and cyclase first acting separately and then simultaneously, on the evolutionary dynamics of the protocell population. For these simulations we assume there are sufficient monomers in the system by taking *S*_*max*_ = 50*V*. We also assume that 8-mer concentration remains fixed for all the simulations. We do these simulations for *V*_*T*_ = 100 and for *N* = 400 protocells, with one circular ssRNA per cell initially, whose replication rate is assigned randomly according to Eq. S1. We found that a replicase alone can never sustain ribozymes in the population for any value of the degradation rate and replicase creation probability. However, for a fixed creation probability, a cyclase alone, can sustain the population below a critical value of degradation rate. This happens because a replicase cannot create new circular templates which it needs to act on to be effective, it can only speed up the replication rates of existing circular strands. As a result, for any non-zero value of the degradation rate, the initial population of circular templates eventually die out without being replenished in the absence of a cyclase, leading to elimination of RNA strands from the protocell population. By contrast, a cyclase alone can create new circular templates and therefore can sustain ribozymes in the protocell population via non-enzymatic replication.

We next consider the case where where replicase and cyclase are simultaneously present in a protocell. At each time step, we measure the fraction of protocells containing either only replicase or cyclase or both or none. Fig-2 shows the time plots of those fractions for different values of the creation probabilities of replicase (*P*_*r*_) and cyclase (*P*_*c*_). As evident from Fig-2(A) when *P*_*r*_ *> P*_*c*_ the abundance of protocells with only replicase (R) and with both replicase and cyclase (RC) are maximum and in equal proportions. The reverse is true for *P*_*c*_ *> P*_*r*_ [Fig-2(B)]. However, for *P*_*r*_ *∼P*_*c*_, protocells with both replicase and cyclase (RC) dominates and their abundance is higher than the maximum abundances of protocells with either of these ribozymes in the previous 2 cases [Fig-2(C)]. This indicates the formation of a positive feedback loop between replicase and cyclase. The cyclase saves the system from degradation by creating new circular strands on which the replicase can act to produce new ssRNA while the replicase boosts replication rates thereby increasing the rate of creation of new replicases and cyclases. Highest abundance of RC protocells along the diagonal of Fig-2(D) (i.e. *P*_*r*_ = *P*_*c*_) further hints at the existence of such a synergistic network between the replicase and cyclase.

**Fig. 2.**
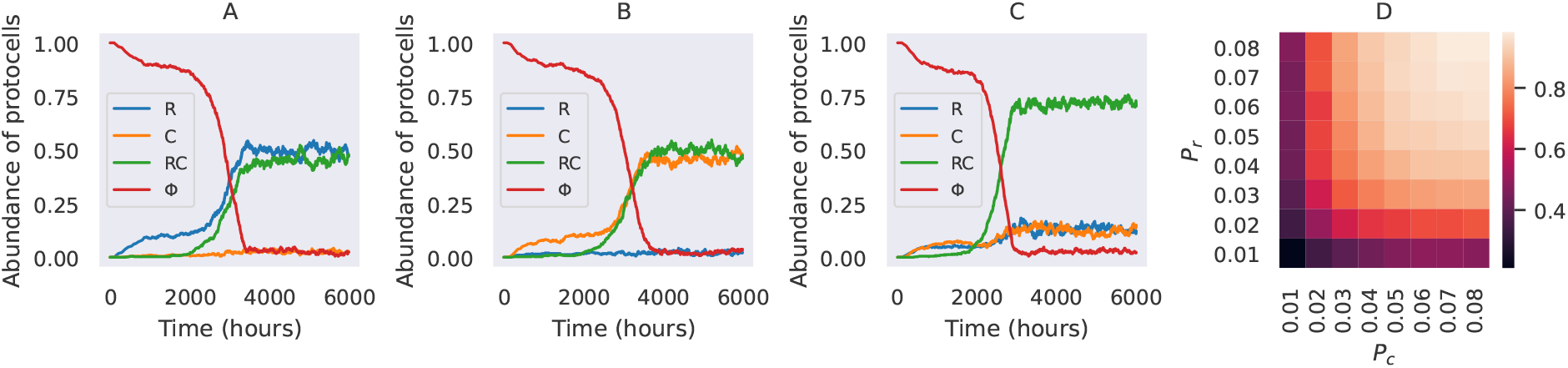
Fractional abundance of protocells containing replicase (R) or/and cyclase (C) or none of them (Φ) for **A:** *P*_*r*_ = 0.05 & *P*_*c*_ = 0.01, **B:** *P*_*r*_ = 0.01 & *P*_*c*_ = 0.05, **C:** *P*_*r*_ = 0.03 & *P*_*c*_ = 0.03 when there are sufficient monomers in the system. **D:** heatmap of the abundance of RC protocells for various value of *P*_*r*_ and *P*_*c*_.

### (b) Role of Nucleotide synthase

If there is a scarcity of monomers in a protocell, the presence of a nucleotide synthase that catalyzes monomer production can be helpful in sustaining growth of RNA strands within a protocell. We model this scenario by taking *S*_*max*_ = 0.8*V* and varying the formation probability of a nucleotide synthase while keeping the formation probabilities for replicase and cyclase fixed. We fixed the strength of nucleotide synthase at *b* = 1. We found that below a threshold value 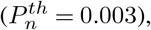, the system dies out since the number of monomers are insufficient to keep the replication process going [Fig-3(A) and (B)]. Above this threshold probability nucleotide synthase ribozymes created inside protocells can provide the extra monomers required at each time step to sustain the replication process. Further a comparison between Fig-3(C) & (D) makes it evident that when nucleotide synthase is created with comparable probabilities as the replicase and cyclase, protocells with all three types of ribozymes dominate the population. When the probability of formation of a nucleotide synthase is much larger compared to that of a cyclase or a replicase, protocells containing either nucleotide synthase alone or together with cyclase or replicase or both cyclase & replicase dominate the population since only protocells containing nucleotide synthase can produce enough monomers necessary for new RNA strand formation, some of which may give rise to replicase and/or cyclase.

**Fig. 3.**
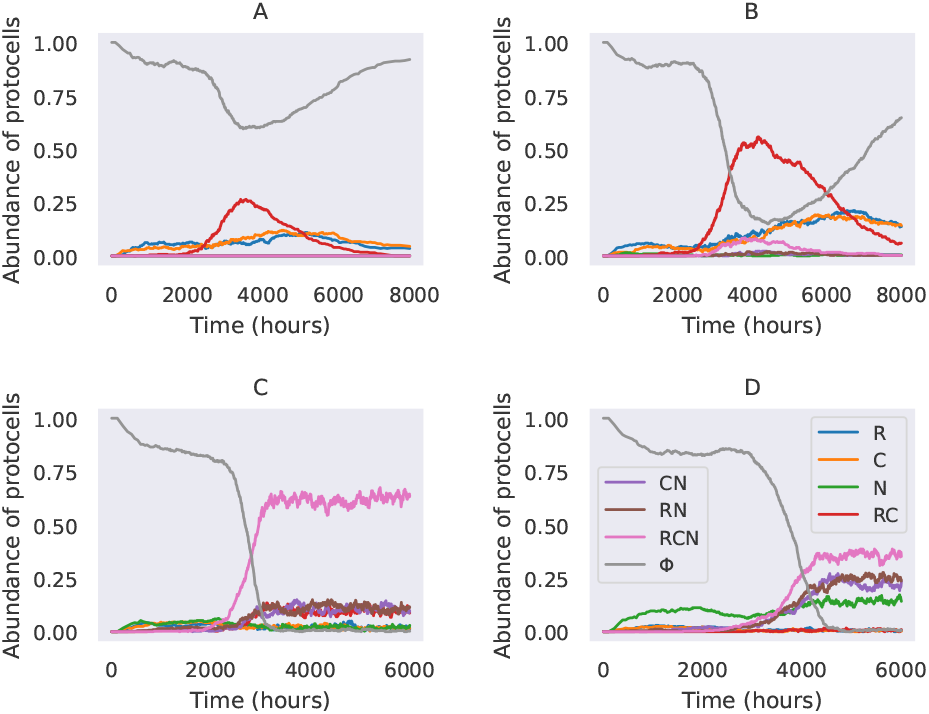
Fractional abundance of protocells containing different types of ribozymes (replicase: R, cyclase: C, nucleotide synthase: N, none: Φ) for **A:** *P*_*r*_ = *P*_*c*_ = 0.03 & *P*_*n*_ = 0.0, **B:** *P*_*r*_ = *P*_*c*_ = 0.03 & *P*_*n*_ = 0.002, **C:** *P*_*r*_ = *P*_*c*_ = 0.03 & *P*_*n*_ = 0.03 and **D:** *P*_*r*_ = *P*_*c*_ = 0.015 & *P*_*n*_ = 0.06, when there is a scarcity of monomers (*S*_*max*_ = 0.8*V*_*T*_, *b* = 1).

### (c) Effect of increasing the threshold volume of proto-cells

The threshold volume of a protocell was so far arbitrarily fixed (*V*_*T*_ = 100) in simulations described in previous sections. The process of membrane transfer from non-lipophilic to lipophilic protocells (containing RNA strands), facilitated by a dipeptide that binds to the membrane [25], can lead to increase in *V*_*T*_ via physical processes only. A peptidyl transferase enzyme [40] can ligate free amino acids to form the peptides needed to initiate this process thereby making it possible for protocells with peptidyl transferase to increase their threshold volume. Increasing *V*_*T*_ has the advantage of ensuring proliferation of protocells with multiple distinct ribozymes for *lower* ribozyme formation probabilities. We obtained the same equilibrium abundances of protocells containing ribozymes by decreasing the probabilities as *P ∝* 1*/V*_*T*_ (see Fig-S7(A-B)). Nevertheless, the advantage of increasing *V*_*T*_ is offset by the decreased likelihood of ribozyme production due to the 1*/V*_*T*_ dependence of the ribozyme catalysed rates. For a particular value of degradation rate there will be a limit up to which the threshold volume can be increased (and simultaneously the formation probabilities can be decreased), progressively making it more difficult for ribozymes to form via enzymatic processes before existing ribozymes are degraded (see Fig-S7(C)). However, for a fixed set of values of the formation probabilities, a protocell with peptidyl transferase is more likely to acquire a selective advantage because of its higher threshold volume. The number of RNA molecules inside such protocells will keep growing beyond the threshold allowed for protocells lacking that ribozyme. This increases the chances of creation of more ribozymes of all types inside them, including peptidyl transferase which will increase the threshold volume even further. This explains the rise in average threshold volume of protocells with time [Fig-4(A)] and the higher value of the relative abundance of cells with all four ribozymes compared to the cases when amino acids and peptidyl-transferase were absent. A comparison between Fig-4 (B),(C) and for the case when *P*_*r*_ = *P*_*c*_ = *P*_*n*_ *> P*_*p*_ makes it clear that protocells with all 4 types of ribozymes are most abundant in the population when all ribozymes are created with comparable probabilities, thereby hinting at the existence of a synergistic network between them. Fig-S8 shows the peak of the distribution of total number of ribozymes & non-catalytic open-ended strands in dividing cells is more than twice as high as those in cells which are getting eliminated by cellular competition. Fig-S9 shows the distribution of the relative abundance of each type of ribozyme in dividing cells.

**Fig. 4.**
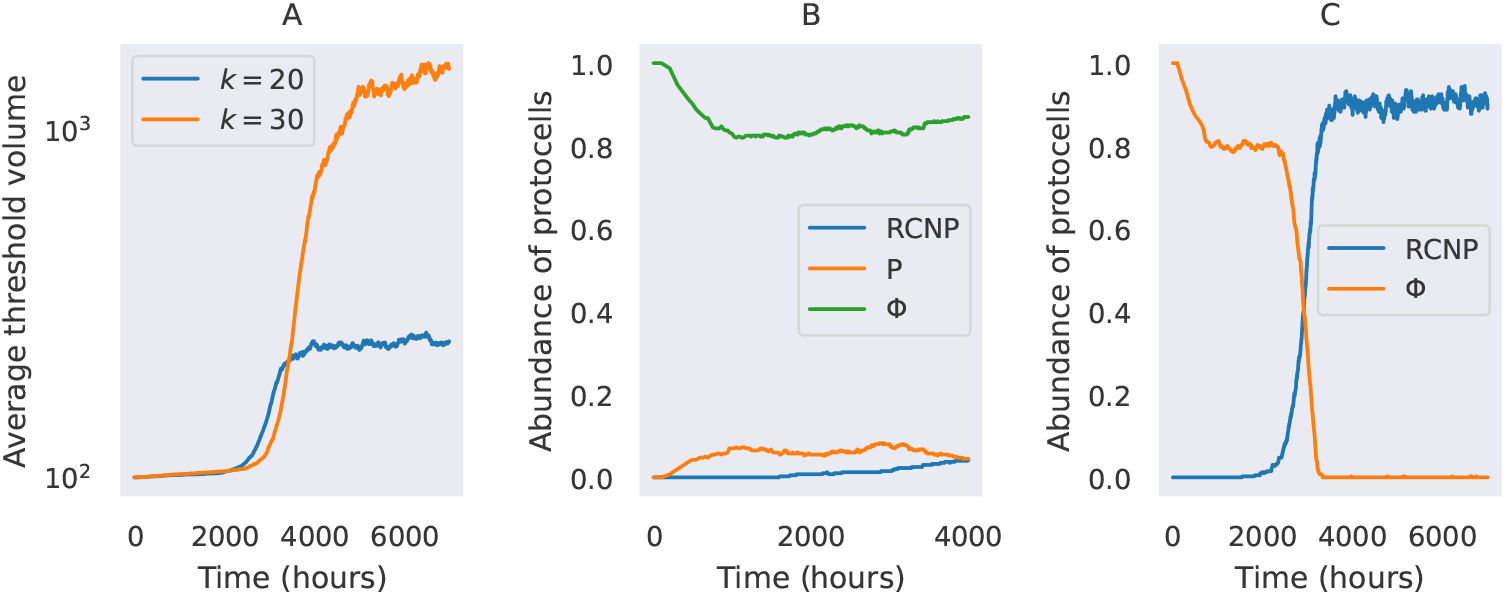
Presence of amino acids and creation of peptidyl-transferase. **A:** Average threshold volume of protocells vs time for 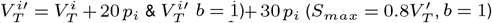 (y axis is in logscale). Fractional abundance of protocells containing ribozymes (RCNP: replicase + cyclase + nucleotide synthase + peptidyl transferase, Φ: none) vs time for **B:** *P*_*r*_ = *P*_*c*_ = *P*_*n*_ = 0.02 & *P*_*p*_ = 0.06 and **C:** *P*_*r*_ = *P*_*c*_ = *P*_*n*_ = *P*_*p*_ = 0.03, when 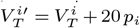.

## 4. Discussion and conclusion

The plausibility of life based on RNA requires RNA encapsulated within lipid vesicles to not just replicate but also acquire a variety of functions that will ensure that the protocell turns into a self-sustaining replicator with heritable traits. A population of protocells can then undergo Darwinian evolution in a manner that allows for the emergence and eventual proliferation of more improved protocell variants. Emergence of functionally diverse ribozymes within a protocell requires rapid replication and we show here how the rolling circle mechanism can be harnessed to ensure exponential growth in the number of long RNA strands inside a protocell. A large number of such strands that can fold into complex secondary structures enhance the likelihood of chance emergence of functional ribozymes. We suggest a hierarchy for the emergence of different ribozymes that would be most beneficial for preferential selection of protocells with more functionally diverse ribozymes during the course of evolution. By varying the formation probability of different ribozymes, it is possible to uncover the conditions under which evolution of the protocell population leads to proliferation of those protocells that contain larger numbers of functionally distinct ribozymes. During this evolutionary process, we highlight the key role that may have been played by lipids as well as small catalytic peptides [41, 42]. The formation of the latter may have been facilitated by the chance emergence of peptidyl transferase ribozymes.

We carried out simulations for a fixed length of RNA strands (200-mers). However, Deamer and collaborators obtained a distribution of circular ssRNA molecules of various lengths (60 to 1100 nt) with the average being *∼*200 nt [39]. Strands with smaller lengths will replicate faster than 200-mers. However the ribozymes like replicase, peptidyl-transferase have lengths *∼*200 nucleotides [6, 40] and nucleotide synthase can also have lengths ranging from 127 to 272 nucleotides [43]. Therefore, only circular strands with lengths *∼*200 nucleotides can produce such functional ribozymes through the rolling circle replication mechanism and protocells containing such ribozymes will grow in size and replicate faster compared to the ones with shorter strands that are non-functional. The former will therefore soon dominate the population. The folded open-ended ssRNA molecules inside protocells that are not ribozymes but have greater abundance than the ribozymes can also ligate with each other to form even longer RNA strands that can become functional.

The nature of evolution discussed here is not strictly evolution in the Darwinian sense where sequence-encoded, heritable, phenotypic traits are selected. In the primordial epoch under study, evolution is driven primarily by biophysical processes that make protocells with larger number of RNA sequences grow at a faster rate. Initially, selection does not distinguish between the nature of the RNA sequence fragments inside the protocells. Eventually however, prootocells with functional ribozymes are selected, not because selection acts on such catalytic phenotypes, but because protocells containing such ribozymes produce more RNA sequences and therefore grow at a much faster rate than protocells lacking those ribozymes. Nevertheless, such ribozymes should not be considered as a heritable trait until they are genetically encoded. Hence, for functional diversity of ribozymes in protocells to be maintained across generations, ribozymes will have to be produced with similar probabilities through the error-prone replication process every generation until the emergence of template specificity. To underscore this point, we carried out simulations where the ribozyme creation probabilities are reduced ten-fold after the fraction of protocells with all four ribozymes increases to 0.5. Even in such a scenario, we find that subsequent evolution can no longer sustain protocells containing these ribozymes (see Fig-S10(A)). We argue that the low fidelity of the replication process was initially advantageous in creating a diversity of complex, secondary structures from a single (or a few) template(s) that increased the chances of emergence of distinct functional ribozymes prior to their encoding in the template sequence. An early appearance of such genetic encoding might occur through the emergence of template specificity that allows higher-fidelity, template-directed, catalysed replication of specific ribozymes. Such templates could be thought of as quasi-genes and a collection of distinct quasi-genes, each producing a specific type of ribozyme, might have been a precursor of RNA genomes where different segments encode different functions. We modeled the emergence of template specificity and higher replication fidelity by biasing the replicase catalysed replication towards creating more replicases. Other ribozymes could still be created during the enzymatic replication process, albeit with a much lower probability. As expected, this leads to significant reduction in the fraction of protocells containing all four ribozymes but increase in the fraction of protocells containing the replicase (see Fig-S10(B)). Eventually, the transition to Darwinian evolution will occur only through genetic encoding of heritable traits. This would require not just improved replication accuracy, brought about by the emergence of the replicase ribozyme, but also functional differentiation of replicases to ensure replication of not just catalytic RNA but also of the templates encoding such traits. Takeuchi *et al* have shown how the division of labour between the template and catalyst [44, 45] could have evolved through conflicting multi-level evolution [45] which induce the breaking of symmetry between the strands. The strand that looses its catalytic function and reduce its copy number inside a protocell eventually becomes the genome encoding for functional molecules.

Any pathway that leads to the origin of life needs to not only explain the appearance and proliferation of the earliest ribozymes but also explain the eventual emergence of a genome encoding these functional molecules. Even though our model does not address the latter phenomenon, which resulted in the encoding of heritable traits, it provides a plausible scenario for the origin and proliferation of protocells containing a functionally diverse set of ribozymes. Our work reinforces the importance of a mixed RNA-lipid-protein world in explaining the evolution of primitive cells. We hope our model will motivate experimental investigations on efficacy of the rolling circle mechanism of replication during this early epoch of protocell evolution.

## Acknowledgments

We thank Julien Derr and Paul G. Higgs for interesting discussions and valuable feedback. S. Roy is supported by an INSPIRE graduate fellowship given by SERB, Government of India.

