## Supplementary material for "Evolution towards increasing complexity through functional diversification in a protocell model of the RNA world"

### Comparison between different replication processes

#### Template-directed primer extension.

In this section we estimate the efficiency of template-directed primer extension process in sustaining exponential growth of the number of RNA strands for various dry phase temperatures. We start our simulation at the beginning of a semi-wet phase starting with the equilibrium concentration of 40-mer templates (68 nM) in a volume of $1\mu m^3$ i.e. with $\sim 40$ random templates (see calculation below). Primers need to be at least 8 nucleotides long in order to form stable pairs with templates in the semi-wet phase (temperature $\sim 37°C$). Primers after attaching to the template can be extended one monomer at a time during this phase. During the subsequent dry phase, the primer-template pair can be separated to result in distinct open-ended ssRNA. A set of RNA polymers is expected to form an exponential distribution of lengths under simultaneously occurring concatenation and hydrolysis processes [1]. The average length at equilibrium is found to be [2] $L_C = \sqrt{K_{con}/K_{hyd}} = 4$. Therefore we choose a probability distribution of the form $p(L) = A\ exp(-L/L_C)$, so that it gives the average length $L_{avg} = \int_0^\infty Lp(L)dL / \int_0^\infty p(L)dL = L_C$. In case of a fixed total number of monomers ($N_0$), we need to normalize the number distribution ($N_0 \times p(L)$) in a way such that the total number of free monomers and monomers constituting the polymers sums up to $N_0$. Therefore we get the normalized number distribution $N(L) = (N_0/L_C^2)exp(-L/L_C)$. Hence for initial concentration of monomers = 160 mM (well in saturation), the concentration of 8-mer sequences will be, $C(8) = (160/4^2)e^{-8/4}\ mM$. 8-mer sequences have a sequence space of size $4^8$. Therefore the concentration of each distinct 8-mer sequence will be $C(8, distinct) = (160/4^2)e^{-8/4}4^{-8}\ mM = 20\ nM$. We assume the length of templates to be 40 nucleotides. Using ViennaRNA package [3] we find that in a random pool of 40-mers, 85% strands fold into secondary structures and remaining 15% can act as templates. Hence starting from an initial pool of 160 mM monomers, the concentration of 40-mer templates in equilibrium will be $C(40, template) = (160/4^2)e^{-40/4} \times 0.15\ mM = 68\ nM$. A complementary 8-mer primer can attach to each 40-mer template at its $3'$ end and extend towards $3'-5'$ direction by attachment of consecutive monomers. Experimental primer attachment rates for 8-mer primers is $K_8 \sim 4 \times 10^{10}\ M^{-1}h^{-1}$ [4]. We make the attachment or annealing rate dependent on Gibbs free energy of the formed template-primer pair by multiplying it with $(1 - e^{-|E|/RT})$, so that pairs with lower free energies will have higher annealing rates. Hence for a 8-mer concentration of 20 nM, the annealing rate becomes $K_{ann} = 800(1 - e^{-|E|/RT})\ h^{-1}$. After primer attachment, primers are extended according to the experimental rates used in our previous work [5]. In the dry phase following the initial semi-wet phase, the extended primers can dissociate from the templates by melting. An equilibrium is reached between melting and annealing of template primer pairs. In a population consisting of a single sequence and many copy numbers of it's complementary sequence, the bound fraction in equilibrium denotes the relative number of strands which are paired. Given the equilibrium constant of melting and annealing process $K_{eq} = e^{+|E|/RT}$, the bound fraction can be calculated as [6],

$$f(T) = \frac{1 + C_T K_{eq} - \sqrt{1 + 2C_T K_{eq}}}{C_T K_{eq}}$$

where $C_T$ is the total concentration RNA single strands, including those in paired condition. In our case, we use the bound fraction as the probability of a template-primer pair to remain paired in the dry phase. While calculating the probability we take a reference $C_T = 1\ \mu M$. We neglect melting process in the semi-wet phase because of low temperature. Hence in the next semi-wet phase, the template-primer pairs which survive melting in the dry phase can undergo further primer extension and head towards formation of dsRNA. The melted pairs can re-anneal in this phase. But while trying to re-anneal they will face competition from two 8-mer primers (one complementary to the template and the other complementary to the old, partially extended primer) which will try to anneal to them instead. The annealing rate for 8-mer primers are same as before. For calculating the annealing rate between a template and it's former primer, we note that $1\mu M$ concentration of strands means 600 strands per $\mu m^3$ volume (which is our simulation volume). In this volume there is only one former complementary primer for each template. Therefore the concentration of the former primer is $1/600\,\mu M = 10^{-6} \times 600^{-1}\,M$. The annealing rate also depends on the length of primer as $K_{ann} \propto \sqrt{L}$ [7]. Hence the re-annealing rate between a template and it's former primer is $K_{ann}^{former} = (1 - e^{-|E|/RT})\sqrt{L/8} \times K_8 \times 10^{-6} \times 600^{-1} h^{-1} = 67\sqrt{L/8}(1 - e^{-|E|/RT})\,h^{-1}$. This way a separated template-primer pair can re-anneal or they can form two separate template-primer pairs with new 8-mer primers. After that they will again undergo primer extension in this phase. Simulations are repeated for 50 trials, for each of the different values of the dry phase temperature, and for each trial the simulation runs for 20 days (semi-wet phase = 12 hours, dry phase = 12 hours). We measure the likelihood of observing exponential growth of the number of template-primer pairs in the system by counting the number ($N_C$) of times (out of 50 trials) all of the template-primer pairs transform into non-separable dsRNA molecules and thereby preventing further replication of RNA strands. This probability is $(1 - N_c/50)$. Fig-S1 shows that exponential growth is ensured mainly for temperatures $T_{dry} \geq 60^{\circ}\,C$.

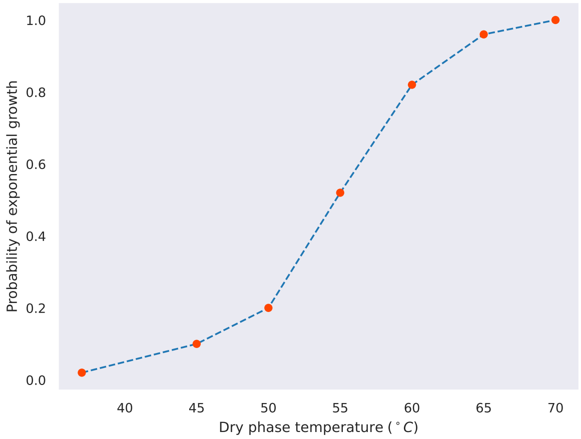


**Fig. S1.** Template-directed primer extension: probability of observing exponential growth of the number of strands vs temperature of the dry phase

### Rolling circle replication.

In this simulation we determine whether the rolling circle replication mechanism can sustain exponential growth of the number of RNA strands. In case of circular templates (circular ssRNA) of length L, 8-mer primers can attach to any of possible L attachment positions and begin primer extension. Hence the primer annealing rate in this case will be $L \times K_{ann} = L \times 800(1 - e^{-|E|/RT})\,h^{-1}$ (assuming the same initial monomer concentration 160 mM as in the previous section) which makes it very high compared to primer extension rates. We therefore make the reasonable assumption that the annealing process is almost instantaneous compared to the primer extension process for this simulation. For comparison with the previous open-ended primer extension process we take the circular templates to be 40 nucleotides in length. As mentioned in the previous section, the initial concentration of 160 mM monomers can give rise to $\sim 40$ (per $\mu m^3$) random templates of length L=40. We assume that with a probability of 0.5, a linear open-ended template can transform into a circular ssRNA template by ligation of its open ends. Therefore we start the simulation with $\sim 20$ random circular ssRNA templates. When the primer on a circular ssRNA becomes full length it becomes a circular dsRNA. Circular dsRNA molecules will undergo further primer extension by rolling circle mechanism. When a circular dsRNA completes one cycle of primer extension by rolling circle method to produce a sequence that is twice the length of the circular template, we assume that the hanging tail detaches itself from the circular dsRNA part. The detached tail will be an open-ended RNA which can fold into secondary structures or act as a linear template with probabilities $(1 - e^{-|E|/RT})$ and $e^{-|E|/RT}$ respectively. As stated earlier, a newly formed linear open-ended template can transform into a circular ssRNA template by ligation of its open ends, with a probability of 0.5. During the rolling circle primer extension process, the hanging tail can also flip at any stage with probability 0.5 thereby reversing the direction of primer extension on the circular template. In simulation we keep track of all circular ssRNA templates, circular dsRNA and open-ended ssRNA molecules. Fig-S2 shows the exponential growth of the number of circular dsRNA and open-ended ssRNA strands.

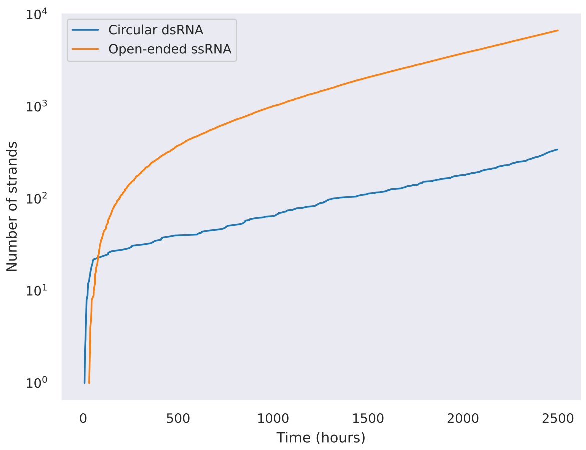


**Fig. S2.** Rolling circle replication: number of circular dsRNA and open-ended ssRNA vs time plot (y-axis is in logarithmic scale)

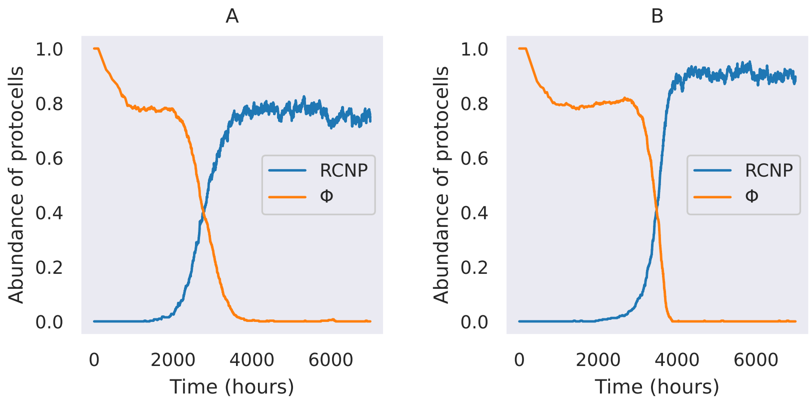


**Fig. S3.** Fractional abundance of protocells containing ribozymes when **A:** the eliminated protocell (labelled by j) in the competition round is selected at random instead of being selected with a probability $\propto (V_i - V_j)$. **B:** a fixed non-enzymatic replication rate $K_{rep} \sim 0.0048\ h^{-1}$ is used all for templates in all protocells.

**Table S1.** Fractional abundance of different secondary structures among the daughter strands created by rolling circle replication from a single circular double-stranded (dsRNA) molecule during 1000 replication events; for different template strand lengths

| Template length | Single hairpin | Double hairpin | Hammerhead | Cloverleaf | More complex |
|---|---|---|---|---|---|
| 50 | 0.706 | 0.28 | 0.001 | 0.011 | 0.0 |
| 60 | 0.446 | 0.489 | 0.007 | 0.058 | 0.0 |
| 70 | 0.364 | 0.499 | 0.028 | 0.107 | 0.002 |
| 80 | 0.339 | 0.489 | 0.048 | 0.12 | 0.004 |
| 90 | 0.153 | 0.505 | 0.103 | 0.218 | 0.021 |
| 100 | 0.138 | 0.421 | 0.107 | 0.294 | 0.04 |

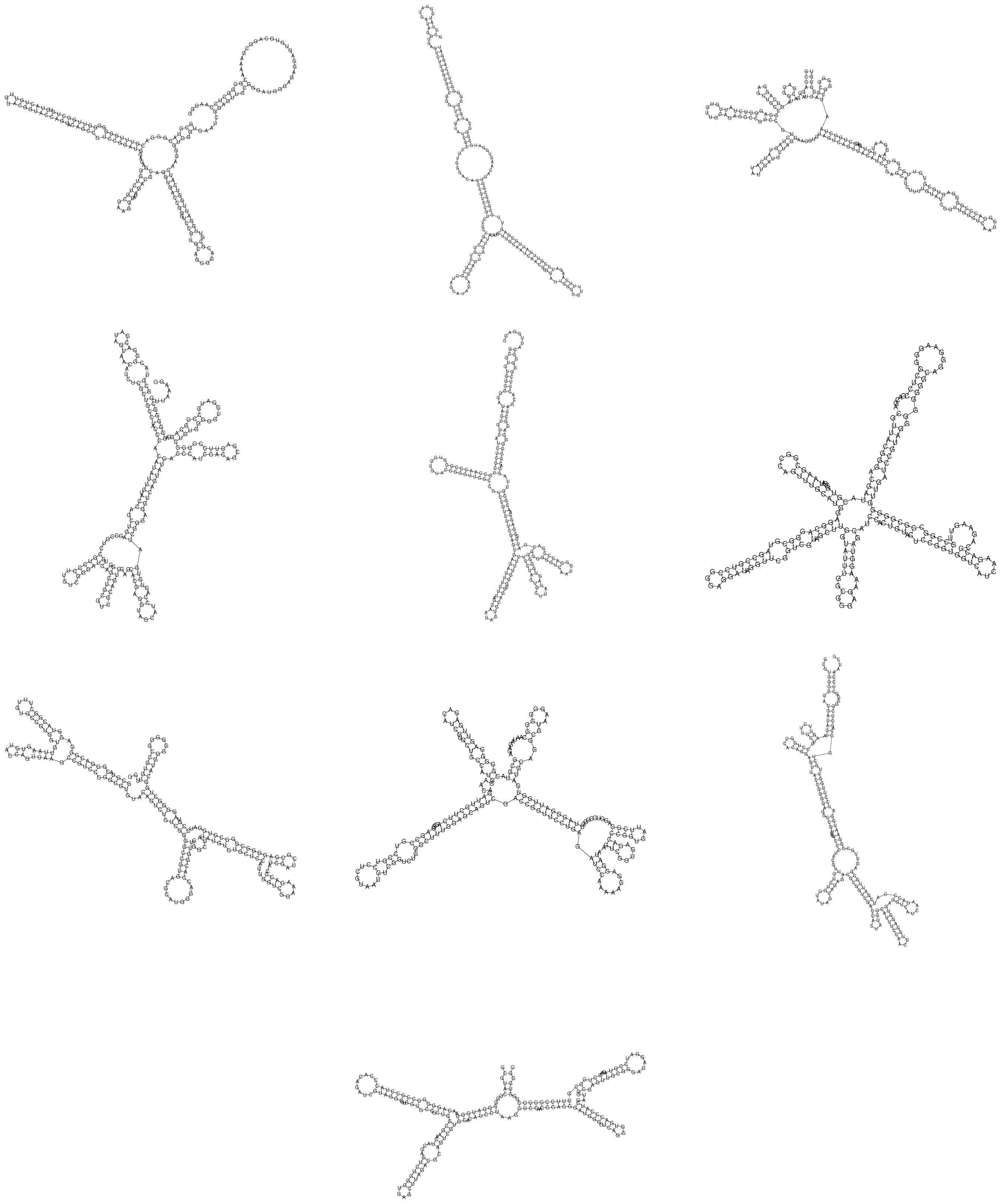

**Fig. S4.** Some diverse secondary structures generated by rolling circle replication from a single circular dsRNA molecule (200 nt)

### Determination of $K_{rep}$

We use sequence-level simulations to determine the non-enzymatic replication rate $K^i_{rep}$ and ascertain its dependence on the length and composition of the sequence. The relative amount of mismatch in a replicated strand is correlated to the sequence of the template. Hence, we first found out the replication rates of 100 random 240-mer sequences and the relative mismatch (number of mismatch/240) for each of the replicated strands using the experimental primer extension rates used in our previous work [5]. We found (by fitting the data) that the replication rate of a sequence depends on its relative mismatch or error as $K_{rep} \propto e^{-b_1(error/L)}$, where $b_1 \sim 2.8$ [Fig-S5(A)]. Further (error/L) follows a normal distribution around a mean of 0.35 with standard deviation 0.0667. To determine the template length dependence of the replication rate, we took each of those templates, kept decreasing their lengths by one nucleotide per step and calculated the replication rate at each step. We found that replication rate depends on the length of a template (for $L \geq 180$ nt) as $K_{rep} \propto K_0 e^{-a_1 L}$, where $a_1 \sim 0.005$ and $K_0 \sim e^{-3.22}\ h^{-1}$ [Fig-S5(B)]. Therefore the non-enzymatic replication rate of a circular template of length L (for $L \geq 180$ nt) is given by,

$$K_{rep} = K_0 e^{-a_1 L - b_1 \times Norm(0.35, 0.0667)}\ h^{-1} \tag{S1}$$

We use this equation to generate the replication rate of a sequence of length L without needing to keep track of the detailed sequence information for each strand in our protocell model. As mentioned in the *Main text* the non-enzymatic replication rates of 200-mer circular ssRNA molecules were chosen from the distribution $K_{rep} = K_0 e^{-a_1 192 - b_1 \times Norm(0.35, 0.0667)}\ h^{-1}$. Nevertheless, protocells containing all four ribozymes continue to dominate (see Fig-S3(B)), even if a fixed non-enzymatic replication rate is used in all protocells.

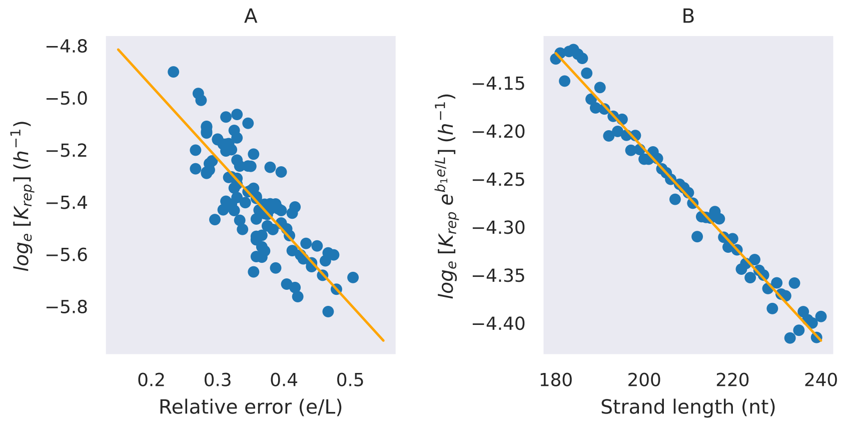


**Fig. S5.** Dependence of non-enzymatic replication rate $K_{rep}$ of circular strands on relative error (e/L) during replication and length of the template strands, as found by using experimental template-directed primer extension rates. **A:** linear fit between logarithm of $K_{rep}$ and (e/L) using random 240-mer templates. **B:** linear fit between logarithm of $K_{rep}$ (excluding the contribution from (e/L)) and length of templates.

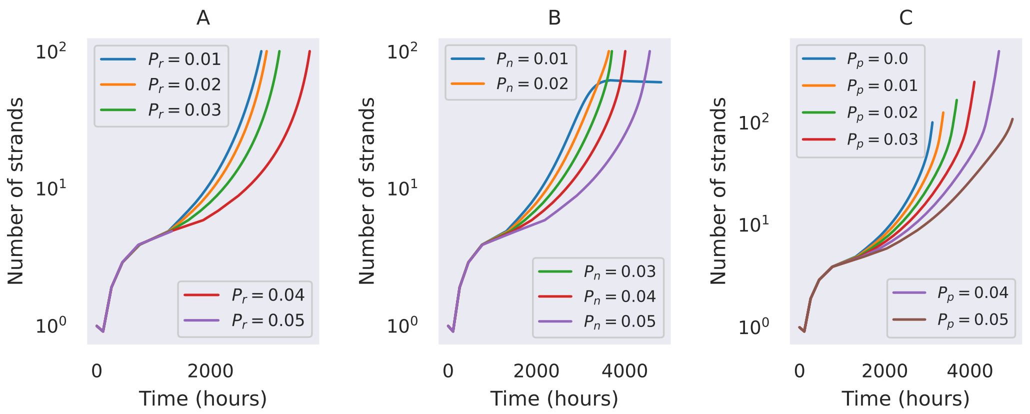


**Fig. S6.** Time evolution of total number of strands inside a single protocell cell. **A:** Variation of replicase creation probability $P_r$ while keeping the value of $P_r + P_c = 0.06$ fixed for the case when monomer availability is not rate-limiting ($P_c$ is the cyclase creation probability). **B:** Variation of nucleotide synthase creation probability ($P_n$) while $P_r + P_c + P_n = 0.09$ and $P_r = P_c$. **C:** Variation of the creation probability of peptidyl transferase ($P_p$) while $P_r + P_c + P_n + P_p = 0.12$ and $P_r = P_c = P_n$.

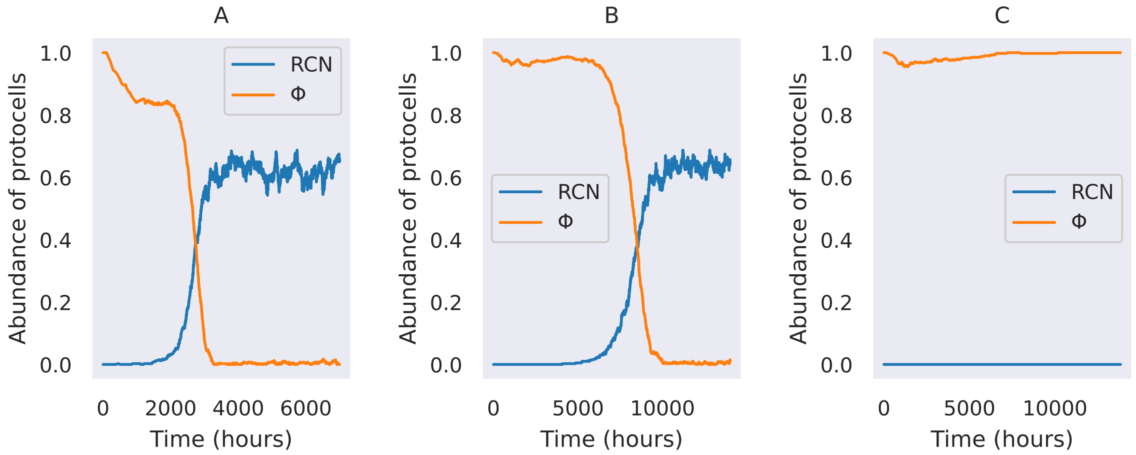


**Fig. S7.** Effect of increasing threshold volume: Fractional abundance of protocells containing ribozymes for diferent threshold volumes when the ribozyme creation probabilities are simultaneously varied as $P \propto 1/V_T$, **A:** $V_T = 100$, $P_r = P_c = P_n = 0.03$; **B:** $V_T = 400$, $P_r = P_c = P_n = 0.0075$; **C:** $V_T = 500$, $P_r = P_c = P_n = 0.006$ $(S = 0.8\,V_T,\ P_p = 0)$.

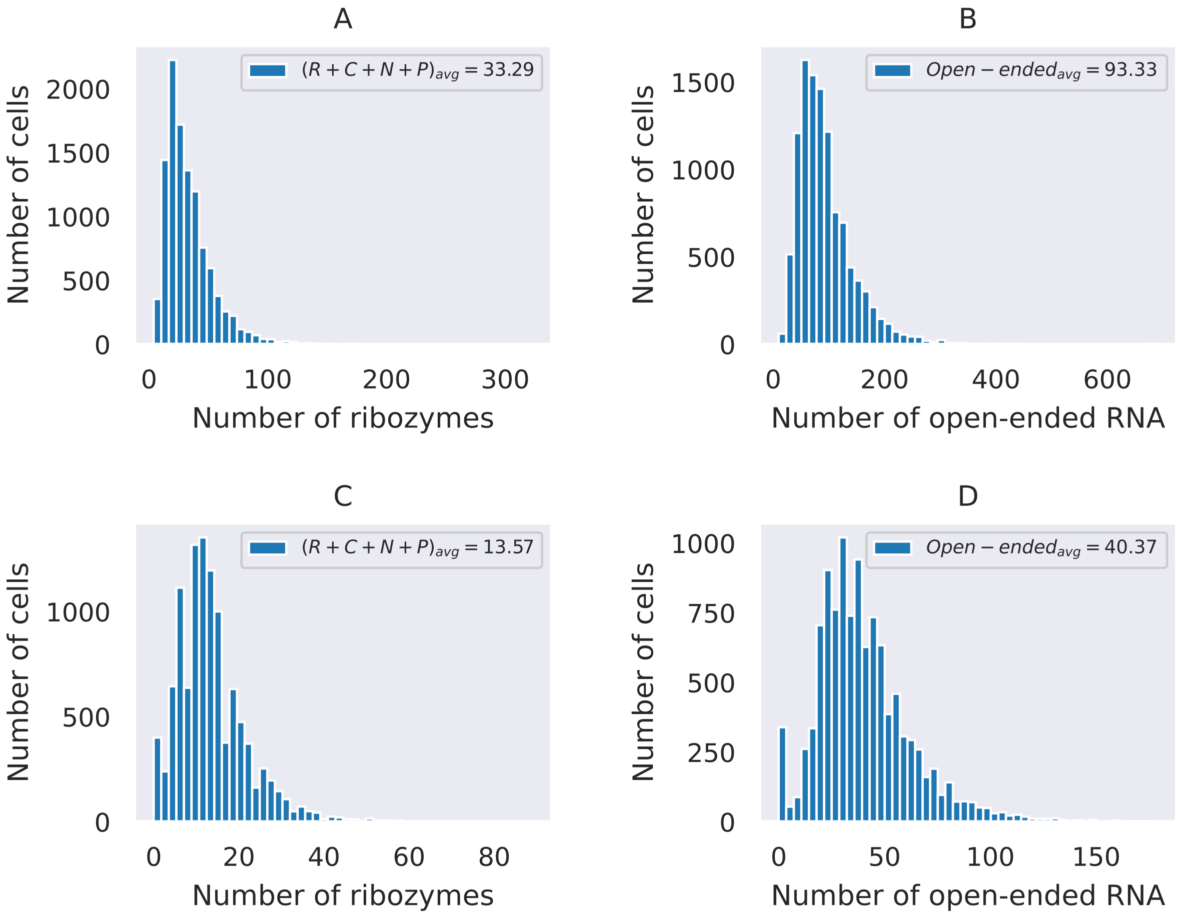


**Fig. S8.** Distribution of total number of ribozymes and non-catalytic open-ended strands respectively for $V_T^{i\prime} = V_T^i + 20p_i$, in **A:** & **B:** dividing protocells. **C:** & **D:** protocells which are getting eliminated.

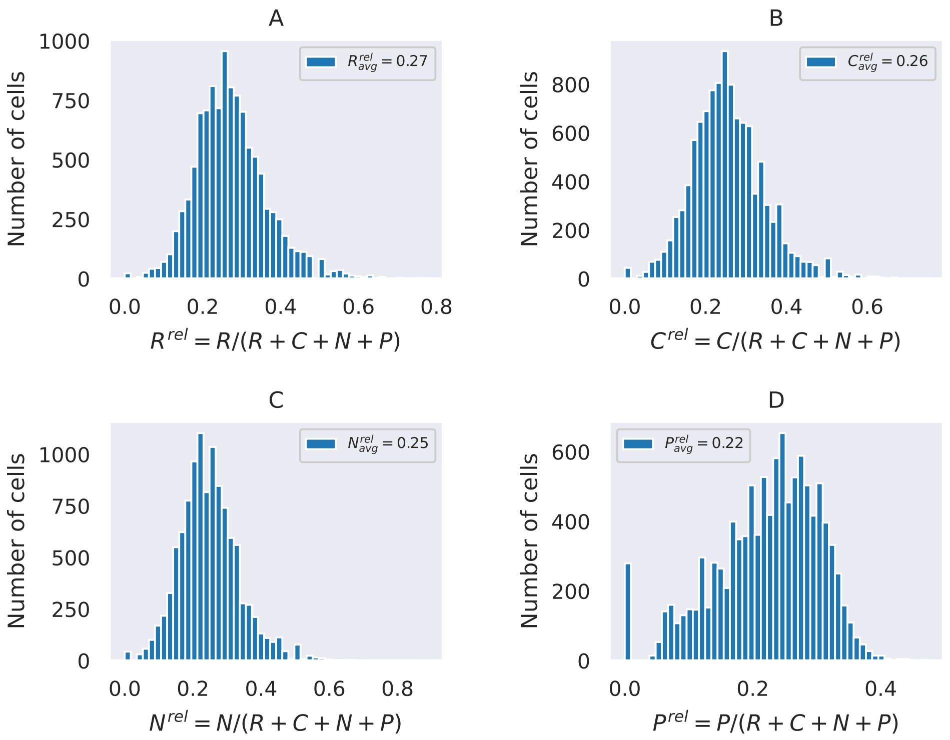


**Fig. S9.** Distribution of the relative abundance of each type of ribozyme (w.r.t. total number of ribozymes) in dividing cells for $V_T^{i\prime} = V_T^i + 20p_i$. **A:** Replicase, **B:** Cyclase, **C:** Nucleotide synthase & **D:** Peptidyl transferase.

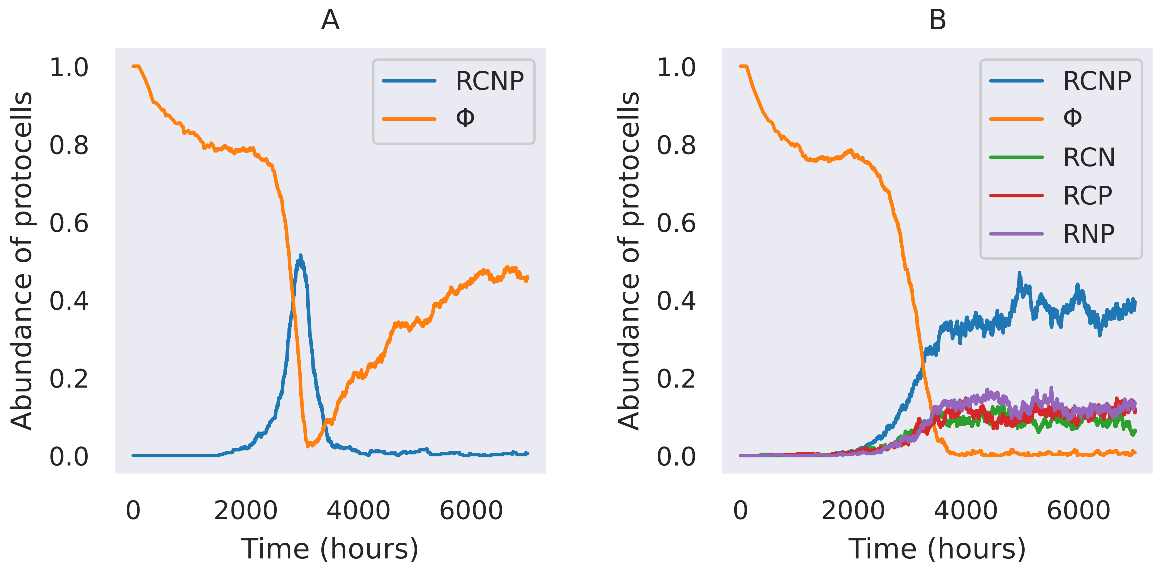


**Fig. S10.** Fractional abundance of protocells containing ribozymes for the case **A:** when we suddenly reduced all ribozymes creation probabilities by 10 fold at the time when abundance of RCNP protocells reaches $50\%$ while not manually changing the $V_T$ value; and the for the case **B:** when we biased the replicase catalyzed replication towards creating more replicases compared to other ribozymes ($P_r^{\prime} = 0.79\,P_r,\ P_c^{\prime} = 0.07\,P_c,\ P_n^{\prime} = 0.07\,P_n,\ P_p^{\prime} = 0.07\,P_p$).

### Pseudocode

**Initial setup:**

Number of cells $N = 400$
$K_{fast} = 0.362\, h^{-1}$
$K_{cyc} = 0.362\, h^{-1}$
Degradation rate $h = 0.0008\, h^{-1}$
$P_r, P_c, P_n, P_p = 0.03, 0.03, 0.03, 0.03$
Create a list of length N for threshold volumes $V_T$ where each element is 100
Create a list of length N for for S where each element is $0.8 \times 100$
Create a list of length N for for $S_{max}$ where each element is $0.8 \times 100$
Create a list named cells, under which there are N empty lists denoting each cell
Fill each cell with 7 empty lists which will contain 7 types RNA strands described in our model
Fill the list for circ ssRNA of each cell with rates randomly assigned as,

$$K_{rep} \sim e^{-3.22-0.005(L-8)-2.8\times rand.norm(0.35,0.0667)}\, h^{-1}$$

Create an empty list 'tot-rate' which will contain the total rates $K^i_{tot}$ for each cell at any time step

**Time evolution:**

Begin loop over time

- Count the total number of nucleotide synthase ($n_o$) across all protocells

  **Determine time step size:**

- Start loop over cells
  - Calculate rate reduction factor $f_i = (S_i + bn_i)/S^i_{max}$
  - Create a list containing the rates for 6 types of reactions:

$$a = \left[\sum_j^{s_i} K^j_{rep} f_i\,,\, \sum_j^{d_i} K^j_{rep} f_i\,,\, K_{fast}\frac{r_i s_i f_i}{V^i_T}\,,\, K_{fast}\frac{r_i d_i f_i}{V^i_T}\,,\, K_{cyc}\frac{c_i l_i}{V^i_T}\,,\, h(s_i + d_i + l_i + r_i + c_i + n_i + p_i)\right]$$

  - Assign $tot - rate_i = sum(a)$
- End loop over cells
- Find $K^{max}_{tot} =$ max(tot-rate)
- Calculate $dt = 1/K^{max}_{tot}$
- Change time $t = t + dt$

  **Reactions:**

- Create a counter variable (g) to count total number of replications
- Start loop over cells
  - If $K^i_{tot} > 0$ and $random.uniform(0,1) < K^i_{tot}/K^{max}_{tot}$
    * Calculate rate reduction factor $f_i = (S_i + bn_i)/S^i_{max}$
    * Create a list containing the rates for 6 types of reactions:

$$a = \left[\sum_j^{s_i} K^j_{rep} f_i\,,\, \sum_j^{d_i} K^j_{rep} f_i\,,\, K_{fast}\frac{r_i s_i f_i}{V^i_T}\,,\, K_{fast}\frac{r_i d_i f_i}{V^i_T}\,,\, K_{cyc}\frac{c_i l_i}{V^i_T}\,,\, h(s_i + d_i + l_i + r_i + c_i + n_i + p_i)\right]$$

    * Choose the type of reaction randomly based the relative propensities ($a/K^i_{tot}$) of the reactions
    * If reaction is circ ssRNA to circ dsRNA
      · Choose a circ ssRNA randomly from the list of circ ssRNA based their relative $K_{rep}$ values

        - Add a $K_{rep}$ to the list of circ dsRNA by multiplying the chosen $K_{rep}$ value with $e^{-0.005L+0.005(L-8)}$
        - Remove the chosen $K_{rep}$ from the list of circ ssRNA
        - Add +1 to g
    * If reaction is circ dsRNA to open-ended ssRNA
        - Draw a random number $p = random.uniform(0,1)$
        - If $p < P_r$ then add a 0 to the list of replicase
        - If $P_r \geq p < P_r + P_c$ then add a 0 to the list of cyclase
        - If $P_r + P_c \geq p < P_r + P_c + P_n$ then add a 0 to the list of nucleotide synthase
        - If $P_r + P_c + P_n \geq p < P_r + P_c + P_n + P_p$ then add a 0 to the list of peptidyl transferase
        - If $p \geq P_r + P_c + P_n + P_p$ then add a 0 to the list of open-ended ssRNA
        - add +1 to g
    * If reaction is replicase catalyzed circ ssRNA to circ dsRNA
        - Choose a circ ssRNA randomly from the list of circ ssRNA
        - Add a $K_{rep}$ to the list of circ dsRNA by multiplying the chosen $K_{rep}$ value with $e^{-0.005L+0.005(L-8)}$
        - Remove the chosen $K_{rep}$ from the list of circ ssRNA
        - Add +1 to g
    * If reaction is replicase catalyzed circ dsRNA to open-ended ssRNA
        - Draw a random number $p = random.uniform(0,1)$
        - If $p < P_r$ then add a 0 to the list of replicase
        - If $P_r \geq p < P_r + P_c$ then add a 0 to the list of cyclase
        - If $P_r + P_c \geq p < P_r + P_c + P_n$ then add a 0 to the list of nucleotide synthase
        - If $P_r + P_c + P_n \geq p < P_r + P_c + P_n + P_p$ then add a 0 to the list of peptidyl transferase
        - If $p \geq P_r + P_c + P_n + P_p$ then add a 0 to the list of open-ended ssRNA
        - add +1 to g
    * IF reaction is cyclase catalyzed open-ended ssRNA to circ ssRNA
        - Add a random $K_{rep}$ value to the list of circ ssRNA following:

$$K_{rep} \sim e^{-3.22-0.005(L-8)-2.8\times rand.norm(0.35,0.0667)}\ h^{-1}$$

        - Remove a $K_{rep}$ value from the list of circ ssRNA
    * If reaction is degradation
        - Choose which type of RNA strand will lose one member, randomly with probabilities proportional to the number of each type in this cell
        - Go the list of the chosen type and remove one member randomly from that list
    * After reaction if $p_i > 0$
        - Change $V_T^i$ as $V_T^i = 100 + 20p_i$
        - Assign $S_{max}^i = 0.8 \times V_T^i$

- End loop over cells
- Calculate $S_{tot} = \sum_i S_i + (bn_o - g)$
- Start loop over cells
    - Assign $S_i = (S_{tot} V_T^i)/\sum_i V_T^i$
    - If new $S_i < 0$, put $S_i = 0$
    - If new $S_i > S_{max}^i$, put $S_i = S_{max}^i$
- End loop over cells

**Cell division:**

- Create a list of cell indices
- Start loop over cells
  - Count total number of strands $(s_i + d_i + l_i + r_i + c_i + n_i + p_i)$ inside the cell
  - If $(s_i + d_i + l_i + r_i + c_i + n_i + p_i) \geq V_T^i$
    - Create an empty list for weakness
    - Start loop over cells
      - If cell index is not marked 'X' and total number of strands inside it $(s_j + d_j + l_j + r_j + c_j + n_j + p_j)$ is lower than that of the dividing cell, then Assign weakness $w_i = (s_i + d_i + l_i + r_i + c_i + n_i + p_i) - (s_j + d_j + l_j + r_j + c_j + n_j + p_j)$
      - Else assign weakness $w_i = 0$
    - Choose a cell randomly with probabilities equal to the relative weaknesses of the cells
    - Make the chosen cell empty
    - Mark the cell index for both the dividing cell and the eliminated cell 'X'
    - Start loop over each strand of the dividing cell
      - Draw a random number p
      - If $p < 0.5$ move the strand to the empty cell and remove from the present cell
      - Else keep the strand in the present cell
    - End loop over strands
    - Change the $V_T$ of the daughter cells as $V_T^i = 100 + 20p_i$ and $V_T^j = 100 + 20p_j$
- End loop over cells

End loop over time
